# SARS-CoV-2 escapes direct NK cell killing through Nsp1-mediated downregulation of ligands for NKG2D

**DOI:** 10.1101/2022.06.20.496341

**Authors:** Madeline J. Lee, Michelle W. Leong, Arjun Rustagi, Aimee Beck, Leiping Zeng, Susan Holmes, Lei S. Qi, Catherine A. Blish

## Abstract

**Summary:** Natural killer (NK) cells are cytotoxic effector cells that target and lyse virally-infected cells; many viruses therefore encode mechanisms to escape such NK cell killing. Here, we interrogated the ability of SARS-CoV-2 to modulate NK cell recognition and lysis of infected cells. We found that NK cells exhibit poor cytotoxic responses against SARS-CoV-2-infected targets, preferentially killing uninfected bystander cells. We demonstrate that this escape is driven by downregulation of ligands for the activating receptor NKG2D (“NKG2D-L”). Indeed, early in viral infection, prior to NKG2D-L downregulation, NK cells are able to target and kill infected cells; however, this ability is lost as viral proteins are expressed. Finally, we found that SARS-CoV-2 non-structural protein 1 (Nsp1) mediates downregulation of NKG2D-L and that Nsp1 alone is sufficient to confer resistance to NK cell killing. Collectively, our work reveals that SARS-CoV-2 evades NK cell cytotoxicity and describes a mechanism by which this occurs.

**Graphical abstract:** 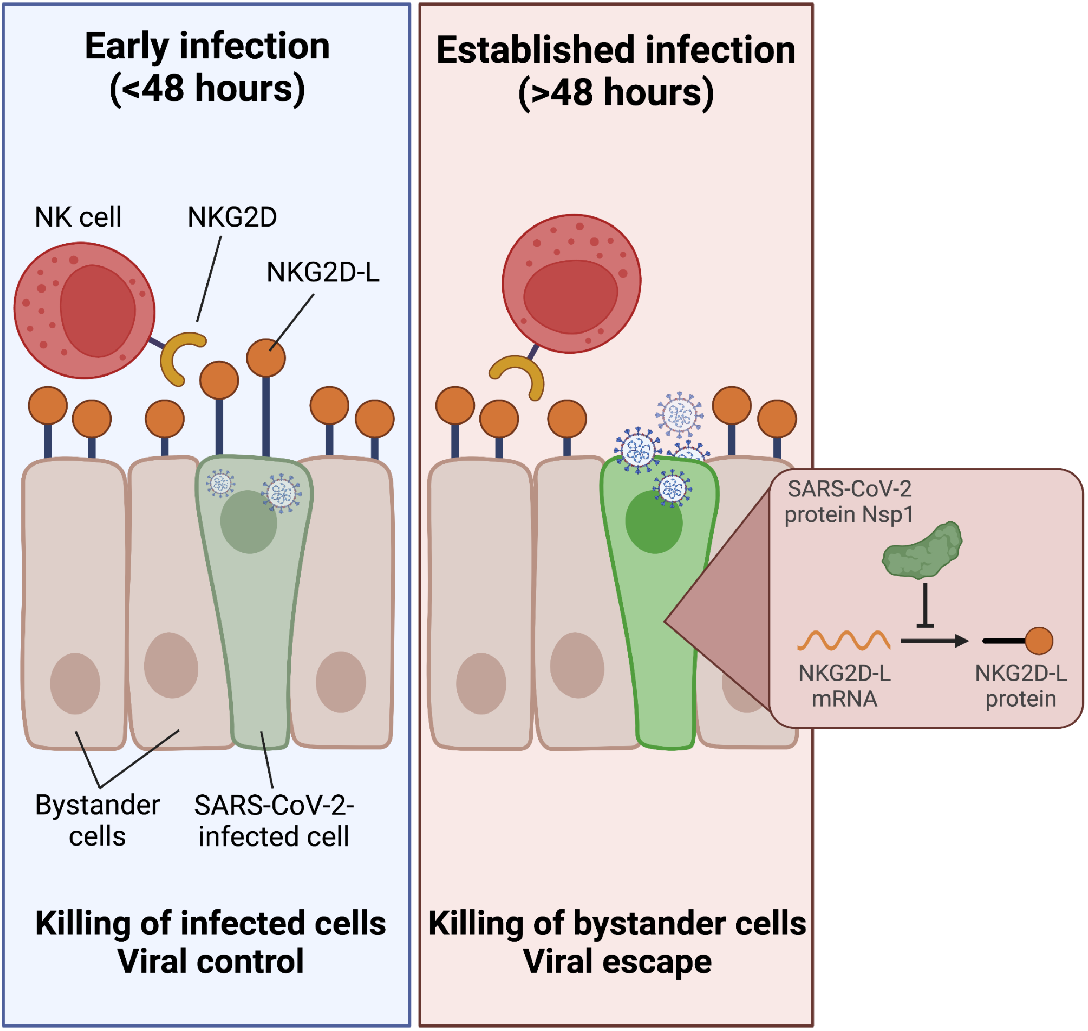

## Introduction

Natural killer (NK) cells are innate lymphocytes that play a critical role in the immune response to viral infection (French and Yokoyama, 2003; Brandstadter and Yang, 2011; Waggoner *et al*., 2016; Björkström, Strunz and Ljunggren, 2022). Since the advent of the COVID-19 pandemic, studies examining the immune response in COVID-19 have noted that NK cells are less abundant in the peripheral blood of severe COVID-19 patients than in healthy donors (Chen *et al*., 2020; Giamarellos-Bourboulis *et al*., 2020; Maucourant *et al*., 2020; Osman *et al*., 2020; Wilk *et al*., 2020, 2021; Zheng *et al*., 2020; Liu *et al*., 2021; Varchetta *et al*., 2021); a concurrent increase in NK cell frequency in the lungs of critically ill patients suggests that peripheral depletion of NK cells may be due to trafficking to the site of infection (Chua *et al*., 2020). Additionally, immune profiling has uncovered significant, severity-associated phenotypic and transcriptional changes in the peripheral NK cells that remain in the blood of COVID-19 patients. In severe COVID-19, peripheral blood NK cells become activated and exhausted (Maucourant *et al*., 2020; Osman *et al*., 2020; Wilk *et al*., 2020, 2021; Zheng *et al*., 2020; Bozzano *et al*., 2021; Krämer *et al*., 2021; Leem *et al*., 2021). They also downregulate surface level expression of the activating receptors NKG2D and DNAM-1, possibly as a consequence of internalization after ligation (Varchetta *et al*., 2021; Wilk *et al*., 2021) and exhibit defects in their ability to respond to tumor target cells and cytokine stimulation compared to NK cells from healthy donors (Osman *et al*., 2020; Zheng *et al*., 2020; Leem *et al*., 2021).

Less is known about how NK cells respond directly to SARS-CoV-2-infected cells, though several studies have demonstrated NK cells can suppress SARS-CoV-2 replication *in vitro* (Krämer et al., 2021; Witkowski et al., 2021; Hammer et al., 2022). Moreover, a recent study discovered that NK cells are able to mount robust antibody-mediated responses against SARS-CoV-2-infected target cells (Fielding *et al*., 2022). However, the mechanisms underlying NK cell responses to SARS-CoV-2-infected cells are not understood. This is particularly important because many viruses employ mechanisms that allow them to evade recognition and killing by NK cells. For example, both HIV-1 and human cytomegalovirus (HCMV) downregulate the ligands for NK cell activating receptors, shielding infected cells from recognition by NK cells (Sutherland *et al*., 2002; Wu *et al*., 2003; Shah *et al*., 2010; Slavuljica, Krmpotić and Jonjić, 2011). In this study, we utilized primary NK cells from healthy donors in conjunction with replication-competent SARS-CoV-2 to create an *in vitro* model system that dissects the NK cell response to SARS-CoV-2-infected cells. We focused on assessing the direct killing of infected target cells in order to better understand how the balance between SARS-CoV-2 recognition and escape contributes to disease. Collectively, our work deeply interrogates the NK cell response to SARS-CoV-2 and provides novel insight into the role of NK cells in COVID-19.

## Results

### Healthy human NK cells respond to SARS-CoV-2 infected target cells *in vitro*

We established a system to explore the NK cell response to SARS-CoV-2 infection using A549-ACE2 cells (Klein *et al*., 2020), which can be recognized and lysed by NK cells and are infectible with SARS-CoV-2. We infected A549-ACE2 cells with SARS-CoV-2/WA1-mNeonGreen (Xie *et al*., 2020) (which replaces ORF7a with mNeonGreen) at a multiplicity of infection (MOI) of 0.5 (Fig. 1A). After 24 hours, approximately 6% of cells fluoresced green, increasing to 50% by 48 hours (Fig. 1A), suggesting that 48 hours is required for robust viral protein expression. To understand how exposure to SARS-CoV-2-infected target cells impacts NK cell phenotype and function, we added NK cells from healthy donors that had been preactivated overnight with IL-2 to the target cells that had been infected for 48 hours (Fig. 1B). This is an important distinction from prior studies that added NK cells immediately after SARS-CoV-2 infection and before the virus-infected cell expresses the full complement of viral proteins (Krämer *et al*., 2021; Witkowski *et al*., 2021; Hammer *et al*., 2022). We then assessed NK cell phenotype and function in NK cells cultured alone vs. those co-cultured with mock-infected cells or with SARS-CoV-2-infected cells for four hours (Fig. 1C,D; Fig. S1A,B).

**Figure 1:**
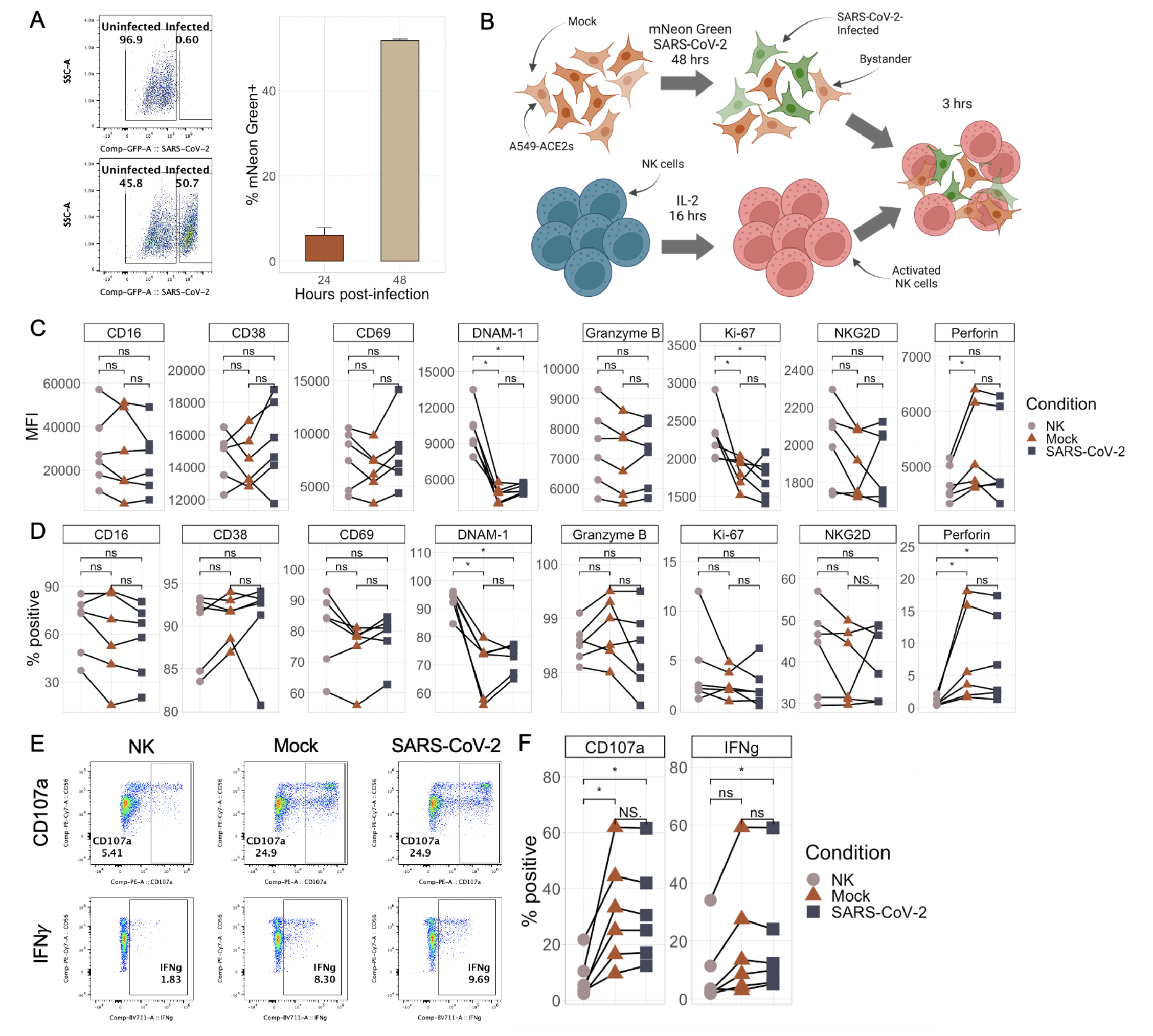
NK cells respond similarly to SARS-CoV-2-infected and mock-infected target cells. A) Representative flow plots (left) and boxplot (right) showing the percentage of mNeonGreen-positive A549-ACE2 cells following infection with either mNeon Green SARS-CoV-2 (MOI 0.5) or media (“mock”) at an MOI of 0.5 for either 24 or 48 hours. Bar plots represent n=4 technical replicates ∓SD values. B) Schematic illustrating the experimental design of NK cell killing assays. C-D) Plots showing the median fluorescence intensity (C) and % of NK cells positive (D) for eight different NK cell markers by flow cytometry upon culture with no targets, mock-infected targets, or SARS-CoV-2-infected targets. E-F) Representative flow plots (E) and quantitations (F) of percentage of NK cells expressing CD107a and IFNγ upon culture with no targets, mock-infected targets, or SARS-CoV-2-infected targets. Significance values were determined using a paired Wilcoxon ranked-sum test with the Bonferroni correction for multiple hypothesis testing.

In response to both mock-infected and SARS-CoV-2-infected A549-ACE2 cells, expression of the activating receptor DNAM-1 and the proliferation marker Ki-67 was significantly downregulated on NK cells (Fig. 1C,D). Similarly, there was an increase in the percentage of perforin-expressing NK cells exposed to both mock-infected and SARS-CoV-2-infected A549-ACE2 cells (though the expression patterns varied between donors). However, in comparing NK cells exposed to infected vs. uninfected cells, there were no SARS-CoV-2 exposure-driven changes in expression of any of the markers assessed, including DNAM-1, CD16, CD38, CD69, NKG2D, Granzyme B, perforin, and Ki-67 (Fig. 1C,D). The same pattern also emerged in functional responses. We observed significant induction of CD107a, a marker of NK cell degranulation and surrogate for cytolytic activity, and IFNγ upon culture with SARS-CoV-2-infected A549-ACE2 cells (Fig. 1E,F); this was not different compared to the response to mock-infected cells.

### SARS-CoV-2-infected cells evade NK cell killing through a cell-intrinsic mechanism

We next assessed whether SARS-CoV-2 modulates the ability of NK cells to kill infected target cells, using the same co-culture system and directly assessing target cell death using a viability dye (Fig. 2A, S2). NK cell co-culture induced significantly more death of uninfected ‘bystander’ cells than of SARS-CoV-2-infected cells in all 17 NK cell donors tested (Fig. 2B). We found no significant difference in the killing of bystander cells compared to mock-infected cells that were never exposed to SARS-CoV-2, indicating that the ability of SARS-CoV-2-infected cells to survive is a cell-intrinsic effect (Fig. 2C). To ensure that these differences were not a result of rapid cell death resulting in cell loss and undercounting of killed SARS-CoV-2-infected cells, we assessed the ratio of infected (mNeonGreen-positive) target cells to uninfected (mNeonGreen-negative) target cells in cultures without NK cells compared to cultures with NK cells, gating only on ‘live’ vs ‘total’ cells. There was no difference in this ratio among all single cells (not live-gated) in the presence and absence of NK cells, suggesting that the cells killed by NK cells are accounted for in our live-dead gating (Fig. 2D). Furthermore, as expected, the ratio of mNeonGreen-positive cells to mNeonGreen-negative cells was increased in live-gated cells upon addition of NK cells due to preferential killing of uninfected target cells by NK cells (Fig. 2D). Collectively, these results support a model in which bystander cells are preferentially killed and imply that there is a factor intrinsic to SARS-CoV-2-infected cells that allows them to escape NK cell killing (Fig. 2E).

**Figure 2:**
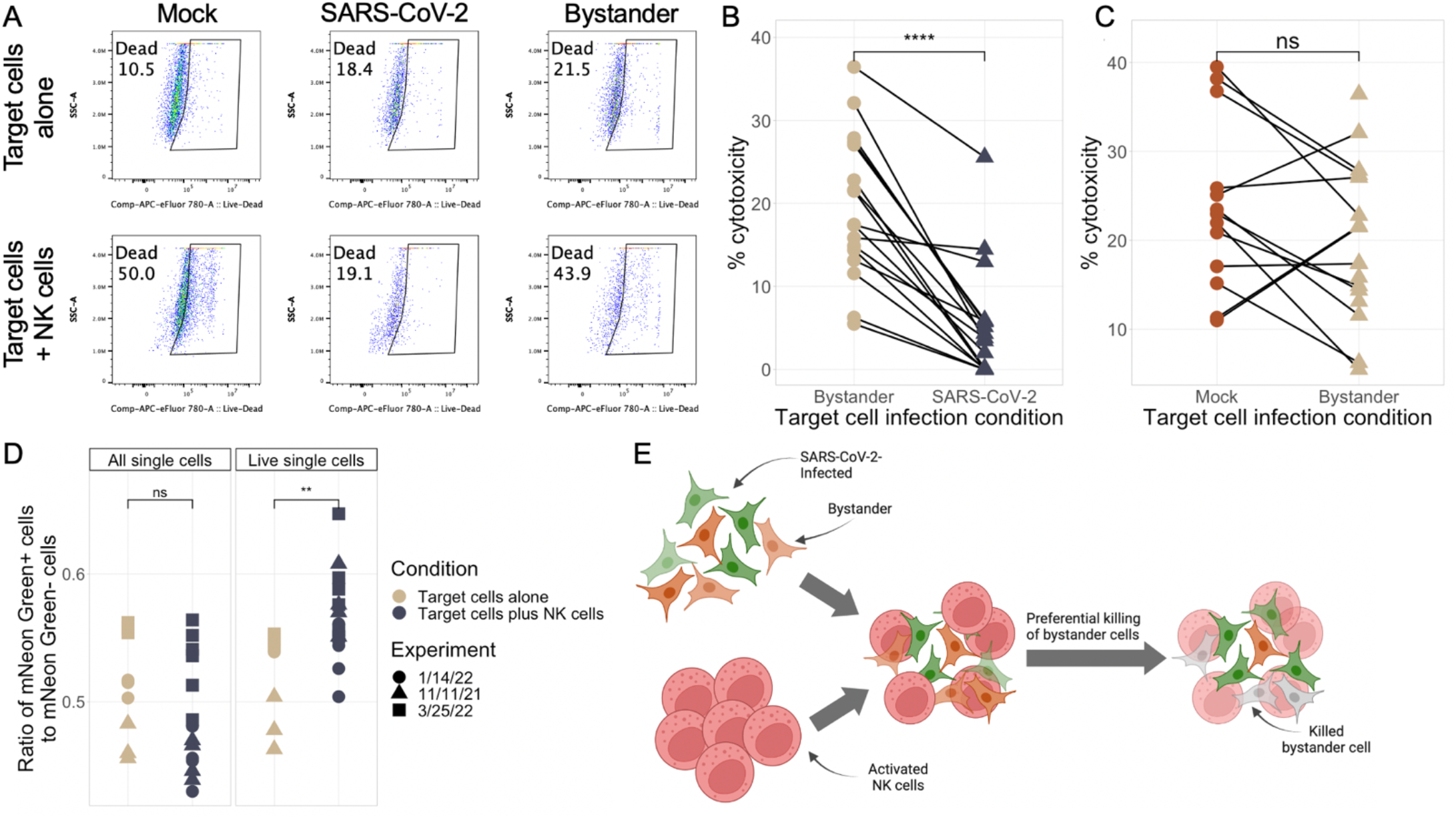
SARS-CoV-2-infected target cells evade NK cell killing through a cell-intrinsic mechanism. A) Representative flow plots showing expression of eFluor 780 viability dye in target cells with NK cells (top) and with NK cells (bottom). B-C) Background-subtracted percentage of A549-ACE2 cell death as measured by eFluor 780 viability dye staining in either infected vs exposed, uninfected cells (B) or mock-infected versus exposed, uninfected cells (C). Background cell death for each experiment and condition was calculated as the average level of death in four wells of the condition of interest. Data are shown from n=4 separate experiments. D) Ratio of mNeon Green positive to mNeonGreen negative A549-ACE2s in the single cell compartment (left) and live single cell compartment (right). E) Schematic illustrating the preferential killing of uninfected bystander cells over infected cells. Significance values were determined using a paired Wilcoxon ranked-sum test with the Bonferroni correction for multiple hypothesis testing.

### SARS-CoV-2 infection modulates expression of ligands involved in NK cell recognition

We next investigated the mechanism by which SARS-CoV-2-infected cells were able to evade recognition and lysis by NK cells. We used flow cytometry to profile the expression of the ligands for various NK cell activating and inhibitory receptors (Björkström, Strunz and Ljunggren, 2022). We grouped antibodies for ligands recognized by the same receptor into a single channel to quantify total ligand density for a given receptor. CD54, CD112/CD155, HLA-ABC, and the ligands for NKG2D were significantly decreased by SARS-CoV-2 infection of A549-ACE2s (Fig. 3A-B, S2A). We also assessed expression of HLA-E and B7-H6 (ligands for NKG2A and NKp30, respectively), which were not detectably expressed in A549-ACE2 cells nor significantly changed by infection (Fig. S2A). While expression of CD112/CD155 (ligands for DNAM-1), CD54 (ligand for LFA-1), and HLA-A/B/C were decreased, the magnitude of these reductions was relatively small. In contrast, the ligands for NKG2D (MICA, MICB, ULBPs 1, 2, 5, and 6; collectively referred to as “NKG2D-L”) were downregulated to a much greater extent in SARS-CoV-2-infected cells compared to uninfected cells and bystander cells (Fig. 3A-B; Fig. S2A). Notably, the downregulation of MHC class I and NKG2D-L would be expected to have opposing effects on the NK cell response to infected cells: downregulation of MHC class I would enhance NK cell recognition of infected targets, while NKG2D-L downregulation could represent a mechanism of NK cell evasion. As we observed a decrease in the ability of NK cells to kill SARS-CoV-2-infected cells, we focused our attention on the downregulation of NKG2D-L as a potential evasion mechanism.

**Figure 3:**
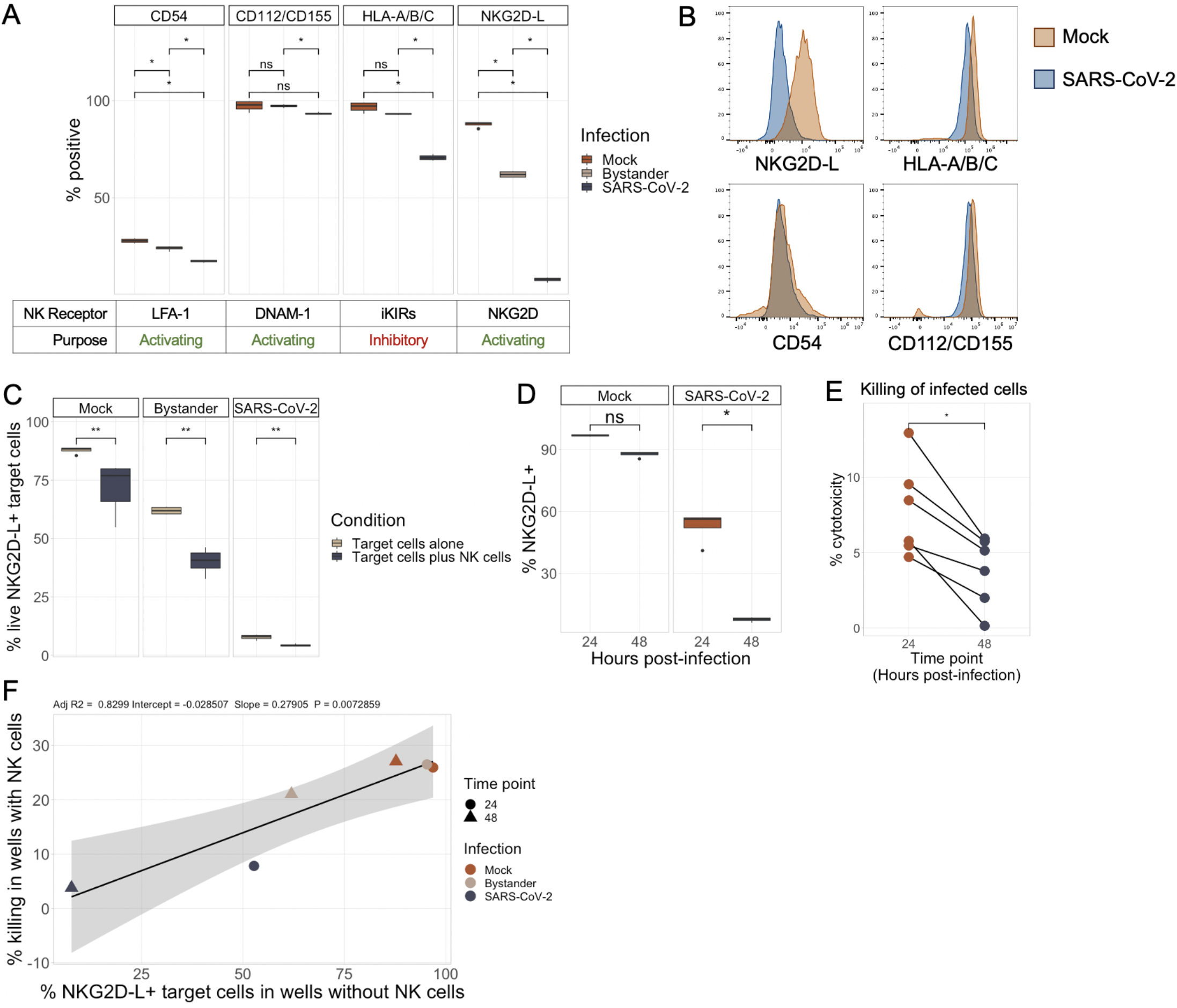
SARS-CoV-2 infection downregulates ligands for the activating receptor NKG2D. A) Boxplots showing the percentage of uninfected, bystander, and SARS-CoV-2-infected A549-ACE2 cells expressing CD54, CD112/CD155, HLA-ABC, and NKG2D-L (combination of MICA, MICB, ULBPs 1,2,5,6). The cognate receptors recognizing each receptor are noted under each pane. 4 technical replicates of each condition were performed. B) Representative histograms of NKG2D-L, HLA-A/B/C, CD54, and CD112/CD155 expression in SARS-CoV-2-infected cells versus uninfected controls. C) Percentage of NKG2D-L-expressing A549-ACE2s in wells containing only target cells compared to wells containing target cells and NK cells. Beginning at 48 hours post-infection, target cells were co-cultured with IL-2-activated NK cells for 3 hours at an E:T ratio of 5:1. D) Percentage of mock-infected or SARS-CoV-2-infected (mNeonGreen+) A549-ACE2 expressing NKG2D-L at 24 and 48 hours post-infection. E) Background-subtracted target cell death of A549-ACE2 infected for either 24 or 48 hours with SARS-CoV-2. Target cells were co-cultured for 3 hours with IL-2-activated NK cells at an E:T ratio of 5:1. F) Correlation between percentage of A549-ACE2s expressing NKG2D-L in target-only wells and background-subtracted target cell death in wells containing NK cells.

### Downregulation of NKG2D-L is associated with inhibition of NK cell killing of SARS-CoV-2 infected cells

To evaluate the association between NKG2D-L expression and killing of SARS-CoV-2-infected cells, we assessed NKG2D-L expression on the cells that survived following co-culture with NK cells. We identified a significant decrease in the frequency of NKG2D-L-expressing target cells in wells containing NK cells at both time points and across all infection conditions, suggesting that NK cells preferentially kill NKG2D-L-expressing targets in both SARS-CoV-2-infected and mock-infected wells (Fig. 3C). We also assessed the kinetics of NKG2D-L expression on infected (mNeonGreen+) A549-ACE2 and found that, while NKG2D-L were downregulated to some extent at 24 hours post-infection compared to uninfected cells, it was not until 48 hours post-infection that we observed almost total loss of these proteins at the surface level (Fig. 3D). We therefore hypothesized that NK cells would kill infected cells more robustly at 24 hours post-infection compared to 48 hours. Indeed, we observed significantly better killing of mNeonGreen-positive target cells at 24 hours post-infection compared to 48 hours (Fig. 3E). Further supporting a model in which downregulation of NKG2D-L allows for evasion of NK cell killing, we identified a strong correlation between the expression of NKG2D-L in target cells and target cell lysis across all time points and infection conditions (Fig. 3F).

### NK cells are able to efficiently kill SARS-CoV-2-infected cells immediately following infection

Other groups have reported that NK cells are able to successfully suppress viral replication in a system where the NK cells are added to a target cell culture immediately after infection with SARS-CoV-2 (Krämer *et al*., 2021; Witkowski *et al*., 2021; Hammer *et al*., 2022). Given our finding that NKG2D-L are not fully downregulated until 48 hours post-infection, we hypothesized that NK cells might be able to kill virus-infected cells in the early stages of infection, but not later. We therefore repeated our killing assay using infected or mock-infected cell cultures at either 0 hours post-infection (similar to prior studies) or 48 hours post-infection. Because the freshly-infected cells had not yet expressed mNeonGreen at the time of analysis (Fig. 4A), we compared total killing of all target cells in SARS-CoV-2-infected wells at 0 and 48 hours and used a MOI of 3 to infect a greater proportion of cells. We found that, as expected, NK cells were able to robustly kill cells that were freshly infected (0 hours) but not those that had been infected for 48 hours (Fig. 4B). Moreover, NK cells were slightly better at killing infected cells compared to mock-infected cells at the 0 hour time point (Fig. 4C), providing additional evidence that NK cells can successfully target infected cells in the very early stages of SARS-CoV-2 infection, as previously reported (Krämer *et al*., 2021; Witkowski *et al*., 2021; Hammer *et al*., 2022). Finally, we conducted a similar analysis of total cell killing at 24 versus 48 hours post-infection. In accordance with our other findings, we observed that NK cells can efficiently kill virus-exposed cells through 24 hours post-infection, but not at 48 hours (Fig. 4D). Thus, our data and other published works collectively suggest that NK cells are capable of suppressing viral replication, but their ability to do so is significantly hampered if the cell has been infected for at least 48 hours. As NK cells appear to home to the lungs during COVID-19 (Liao *et al*., 2020; Huang *et al*., 2021; Brownlie *et al*., 2022), our findings indicate that the timing of NK cell trafficking to the site of infection may impact the efficacy of the NK cell response to SARS-CoV-2 infection.

**Figure 4:**
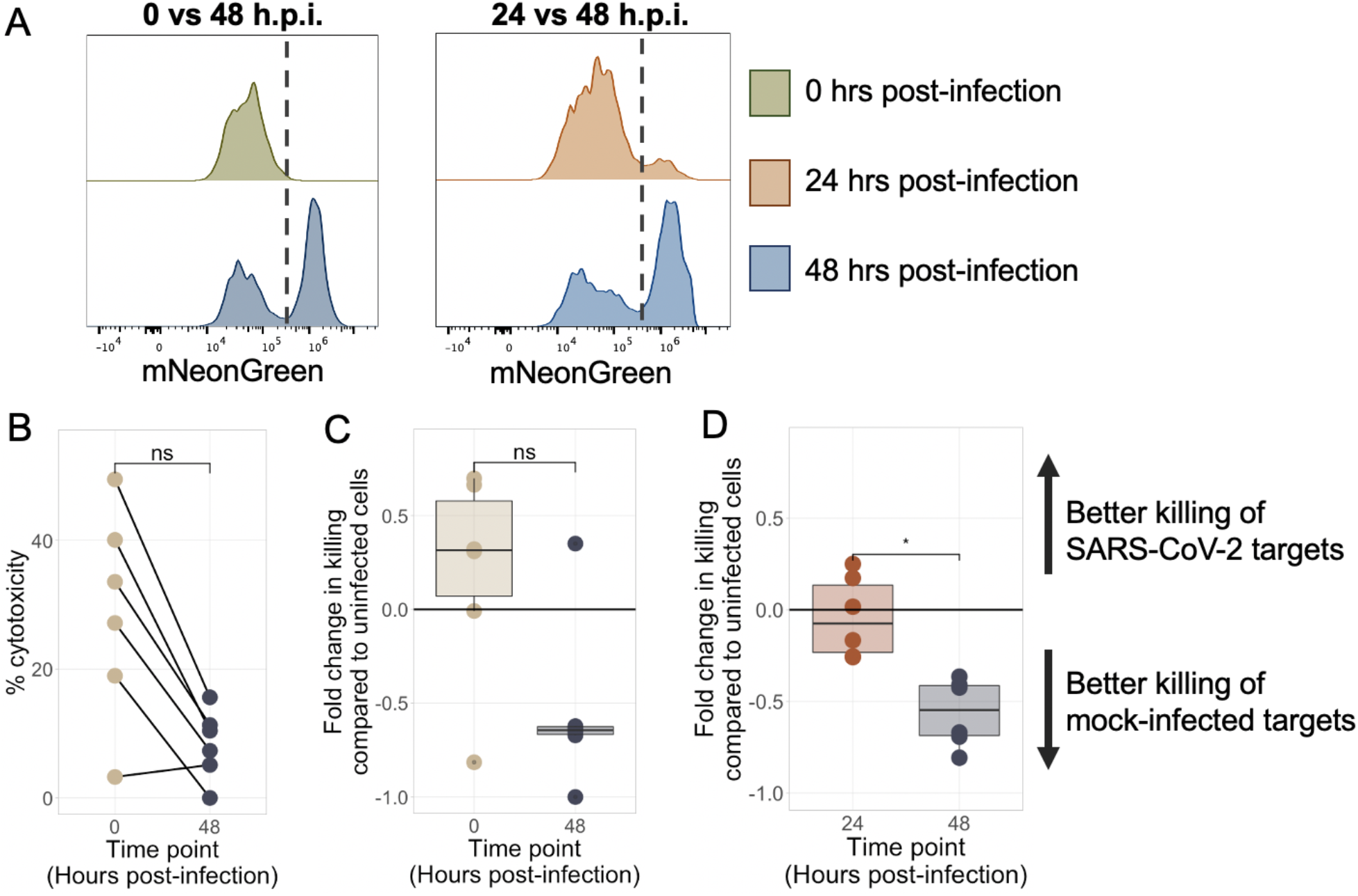
NK cells are able to efficiently kill SARS-CoV-2-infected cells immediately following infection. A) Representative histograms showing expression of mNeonGreen in SARS-CoV-2-exposed A549-ACE2 at 0, 24, or 48 hours post-infection at an MOI of 3 (left) or 0.5 (right). Vertical dashed lines indicate threshold for positive gating. B-D) A549-ACE2 were infected with SARS-CoV-2 for 0, 24, or 48 hours, then co-cultured with IL-2-activated NK cells for 3 hours at an E:T ratio of 5:1. B) Background-subtracted killing of all single A549-ACE2s by NK cells following infection with SARS-CoV-2 for either 0 or 48 hours. C) Fold change in killing of infected target cells compared to mock-infected target cells at 0 and 48 hours post-infection. D) Background-subtracted target cell death of all single A549-ACE2s infected with SARS-CoV-2 at 24 and 48 hours post-infection.

### SARS-CoV-2 protein Nsp1 downregulates ligands for NKG2D

Having identified changes in the protein-level expression of NKG2D-L in SARS-CoV-2-infected cells, we next sought to understand how SARS-CoV-2 mediates this effect. SARS-CoV-2 encodes 29 individual proteins that are broadly classified into three categories: structural, non-structural, and accessory. While the roles of these proteins are still being investigated, many of the non-structural and accessory proteins are known to mediate suppression of antiviral innate immune responses (Mariano *et al*., 2020; Naqvi *et al*., 2020; Raj, 2021; Yadav *et al*., 2021). We therefore transfected each individual SARS-CoV-2 protein, tagged with two Strep Tag domains to allow for easy detection, into A549-ACE2 cells and assessed for their effect on NK cell receptor ligand expression by flow cytometry (Fig. 5A,B). We successfully transfected 25 of the 29 SARS-CoV-2 proteins into A549-ACE2s; we also transfected cells with GFP as a non-viral control (Fig. S4). While several proteins downregulated NKG2D-L, SARS-CoV-2 Non-Structural Protein 1 (Nsp1) had by far the strongest effect (Fig. 5C). Nsp1 also mediated downregulation of MHC class I, but not CD54 or the ligands for DNAM-1 (Fig. 5D,E, Fig. S5). Several other viral proteins, primarily accessory proteins, also downregulated NKG2D-L expression, and some increased expression. However, as Nsp1 had by far the strongest impact on NKG2D-L expression, we chose to move forward with interrogation of this protein.

**Figure 5:**
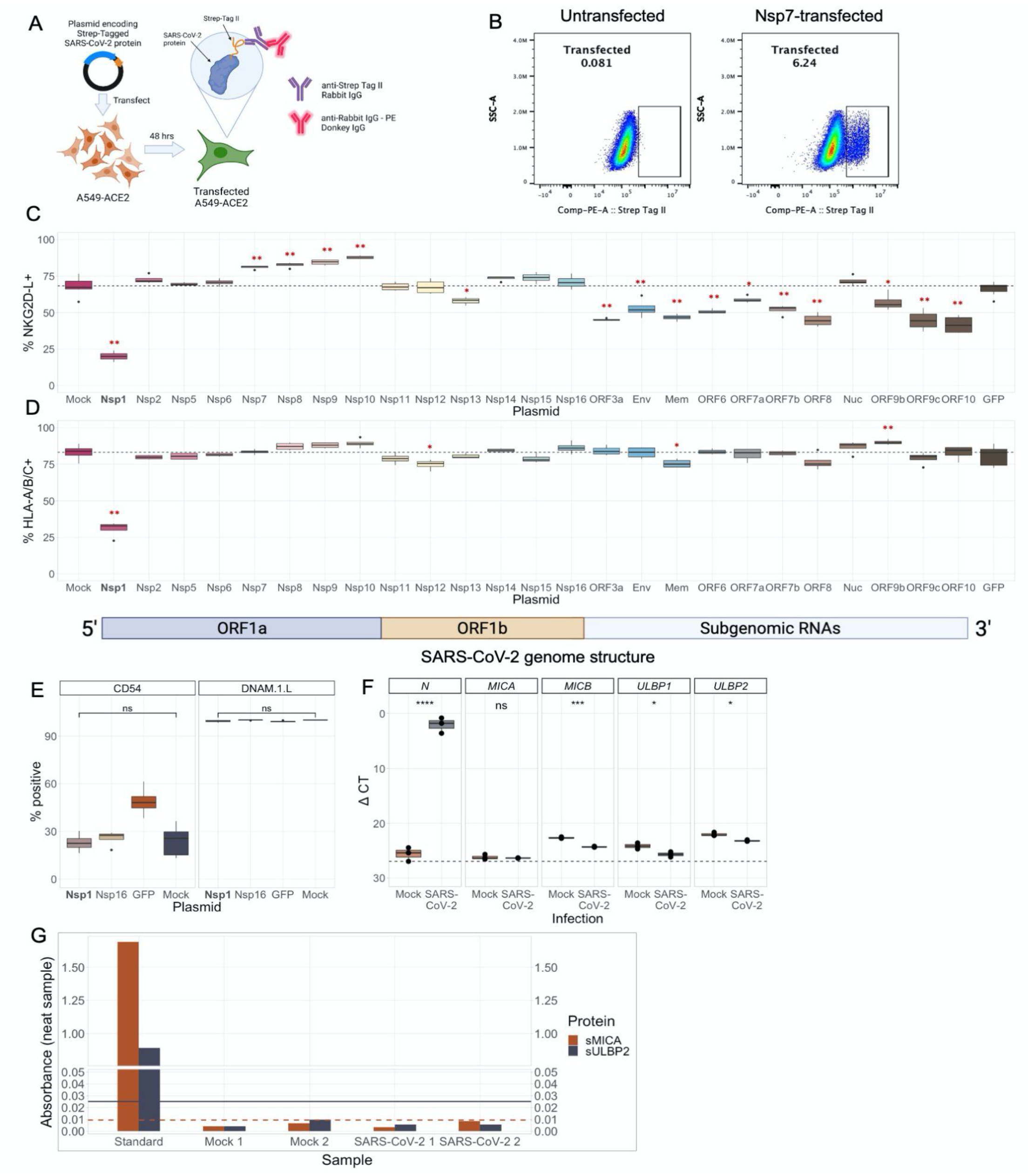
SARS-CoV-2 protein Nsp1 downregulates ligands for NKG2D. A) A schematic illustrating the experimental approach. Plasmids encoding individual SARS-CoV-2 proteins appended with two Strep-Tag domains were transfected into A549-ACE2s. After 48 hours, transfected cells could be detected via flow cytometry using a primary antibody against Strep Tag II and a secondary fluorescent antibody. B) Representative flow plots showing Strep Tag II expression in untransfected and Nsp7-transfected A549-ACE2s. C-D) Percentage out of transfected (Strep Tag II^+^) A549-ACE2s that express NKG2D-L (C) and HLA-A/B/C (D) by flow cytometry at 48 hours post-transfection. 25 SARS-CoV-2 proteins are shown that were successfully transfected into A549-ACE2s, along with GFP as a transfection control. Four technical replicates were performed for each plasmid. Dashed line represents the mean frequency of expression in untransfected (“mock”) cells. Asterisks represent significance in comparison to mock-transfected controls. Plasmids are ordered by location in the SARS-CoV-2 genome and a schematic of the genome structure is shown below Fig. 4D. E) Percentage of A549-ACE2 positive for CD54 or DNAM-1-L (CD112/CD155) at 48 hours after transfection with PBS (mock), Nsp16, GFP, or Nsp1. F) The transcription of ligands for NKG2D are modestly decreased by SARS-CoV-2 infection and these ligands are not secreted. A) Boxplots showing the delta CT values of several genes in mock and SARS-CoV-2-infected A549-ACE2s as measured by RT-qPCR. *N* encodes SARS-CoV-2 nucleoprotein. *MICA, MICB, ULBP1*, and *ULBP2* encode ligands for NKG2D. G) Absorbance values of neat supernatants from mock or SARS-CoV-2-infected cultures at varying dilutions as measured by plate-based ELISAs for soluble MICA (sMICA) and soluble ULBP2 (sUPBP2). Absorbance values were calculated by subtracting absorbance readings taken at 560 nm from those taken at 450 in accordance with the manufacturer’s instructions. Horizontal lines indicate limits of detection (dashed: sMICA; solid: sULBP2). Bar plots represent the means of two technical replicates for each condition.

### Nsp1 post-transcriptionally downregulates NKG2D-L and does not induce shedding

Nsp1, also known as the SARS-CoV-2 leader protein, is the first protein translated when the virus enters a cell and serves as a global translation inhibitor. Nsp1 is highly conserved across coronaviruses as it plays an important role in enhancing pathogenicity by inhibiting the innate immune response (Kamitani *et al*., 2006; Züst *et al*., 2007; Narayanan *et al*., 2008; Min *et al*., 2020; Schubert *et al*., 2020; Vazquez *et al*., 2021). Schubert et al demonstrated that SARS-CoV-2 Nsp1 functions by sterically inhibiting entry of mRNA into the mRNA channel of the 40S ribosomal subunit (Schubert *et al*., 2020). Thus, it is likely that Nsp1 mediates a translational block to reduce surface NKG2D-L expression. Consistent with this model, we observed only a small decrease in transcripts encoding MICB, ULBP-1, and ULBP-2 in infected cells compared to mock-infected cells (Fig. 5F). This difference was quite modest and likely reflects the overall decrease in transcript levels in cells infected with SARS-CoV-2, and is consistent with the idea that NKG2D-L expression is reduced at the post-transcriptional level. We also addressed another possibility, shedding of NKG2D-L from the cell surface, which has been reported for other viruses and in the setting of cancer (Raffaghello *et al*., 2004; Slavuljica, Krmpotić and Jonjić, 2011; Baugh, Khalique and Seymour, 2020), by assessing NKG2D-L levels in the supernatants of mock- and SARS-CoV-2-infected cultures by ELISA. We quantified levels of soluble MICA (sMICA) and soluble ULBP-2 (sULBP2) (Fig. 5G, Fig. S5). We were unable to detect either of these proteins in the supernatants of uninfected or infected cultures, suggesting that secretion of NKG2D ligands is not the mechanism by which NKG2D-L is downregulated by SARS-CoV-2.

### NKG2D-L have a high rate of surface turnover

Although Nsp1 is a global inhibitor of host translation, our data show that it does not equally downregulate all NK cell receptor ligands on A549-ACE2s. We hypothesized that this might be due to differential rates of surface expression turnover across the various ligands, as these proteins are known to have varying levels of stability on the cell surface (Braun *et al*., 1997; Fernández-Messina, Reyburn and Valés-Gómez, 2016; Toledano *et al*., 2018; Yarzabek *et al*., 2018). NKG2D-L in particular is rapidly turned over in order to allow the body a high degree of control over its expression level (Fernández-Messina, Reyburn and Valés-Gómez, 2016; Toledano *et al*., 2018). To validate that inhibition of global translation or trafficking of proteins could have an outside effect on NKG2D-L in comparison with other ligands such as CD54 and DNAM-1 ligands, we treated A549-ACE2s with the protein transport inhibitor Brefeldin A and measured expression of NK cell receptor ligands after 24 or 48 hours (Fig. S6). We observed that Brefeldin A, like Nsp1, had a much larger effect on NKG2D-L than on other ligands, including CD54 and DNAM–1 ligands, supporting a model in which global translation inhibition, such as that mediated by Nsp1, could much more dramatically downregulate NKG2D-L than other surface proteins.

### Nsp1 is sufficient to confer significant resistance to NK cell-mediated killing

Finally, we hypothesized that, if Nsp1 is the key mediator of NKG2D-L downregulation in SARS-CoV-2 infection, transfection with Nsp1 should be sufficient to confer resistance to NK cell killing. To test this hypothesis, we co-cultured activated, healthy NK cells with cells that had been transfected with either Nsp1 or a control plasmid (GFP) and assessed target cell killing by flow cytometry. Indeed, we found that NK cells were significantly more effective at killing GFP-transfected targets compared to Nsp1-transfected targets (Fig. 6A,B; Fig. S7). We also compared killing of Nsp1- and GFP-transfected target cells to killing of cells transfected with another SARS-CoV-2 protein, Nsp14 (Fig. 6C, D); Nsp14 was shown to downregulate NKG2D-L in 293T cells (Fielding *et al*., 2022), although it did not do so in A549-ACE2s in our study (Fig. 5C). Nsp14-transfected cells were killed at similar rates to GFP-transfected cells and were killed significantly more frequently than Nsp1-transfected cells. These data establish that SARS-CoV-2 Nsp1 is sufficient for evasion from NKG2D-L-mediated killing.

**Figure 6:**
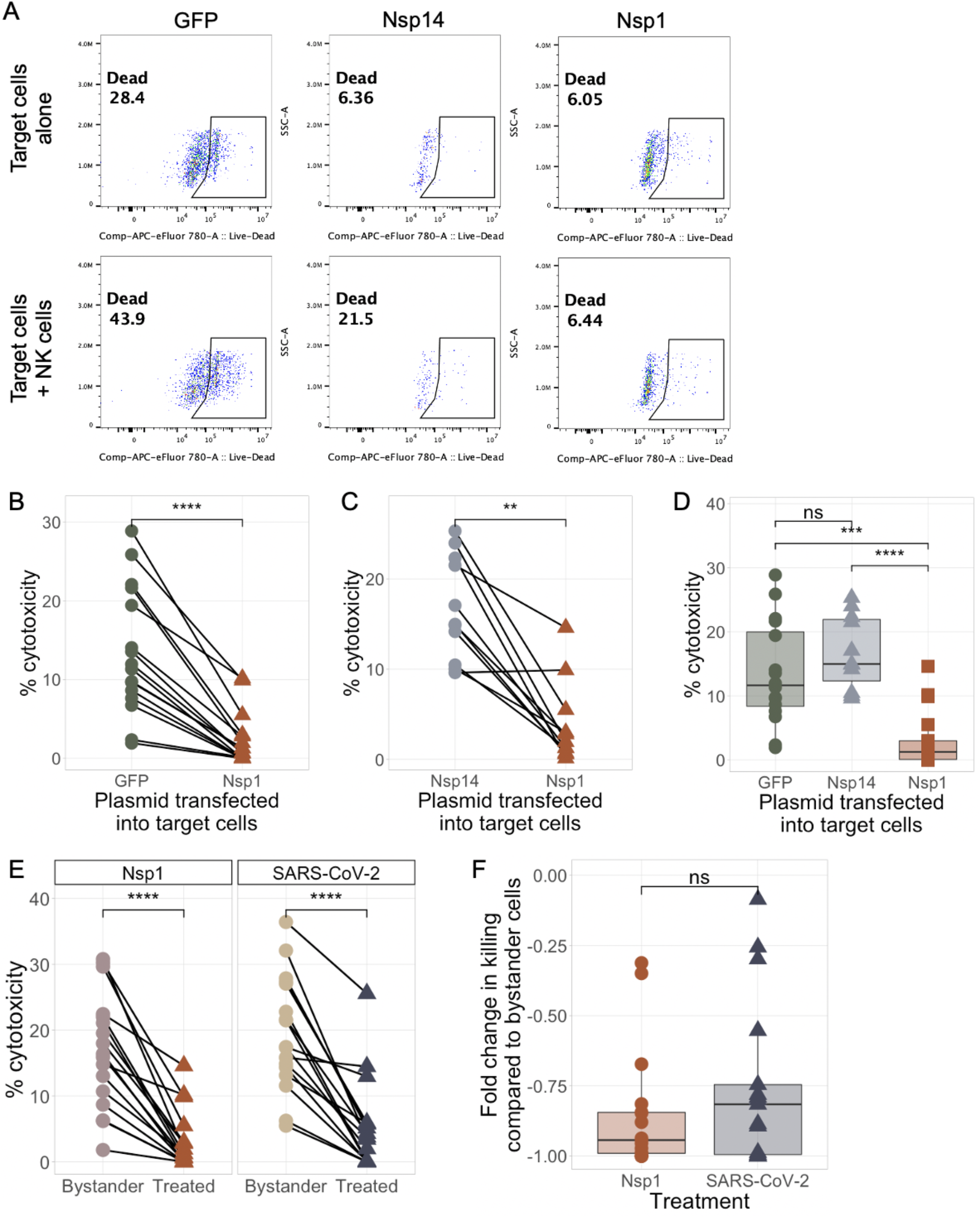
Nsp1 is sufficient to confer significant resistance to NK cell-mediated killing. A) Representative flow plots showing expression of eFluor 780 viability dye in target cells with NK cells (top) and with NK cells (bottom). B-E) Background-subtracted target cell death among cells transfected with either GFP, Nsp1, or Nsp14 and co-cultured with healthy NK cells (E:T = 5:1) for 3 hours. Transfected cells were gated by Strep Tag II expression before % cytotoxicity was determined. Lines in B, C, and E represent individual donors. F) Boxplot showing the fold change in background-subtracted target cell death between bystander (uninfected/untransfected) cells and cells that were positive for either Nsp1 (transfected) or SARS-CoV-2 (infected). Significance values were determined using a paired Wilcoxon ranked-sum test (B, C, E) or unpaired Wilcoxon ranked-sum test (D, F) with the Bonferroni correction for multiple hypothesis testing where necessary.

## Discussion

The role of NK cells in mediating clearance of SARS-CoV-2-infected cells *in vivo* remains unclear. While several studies have demonstrated that NK cells can reduce the levels of SARS-CoV-2 replication *in vitro*, no prior study has directly evaluated killing of SARS-CoV-2 infected cells. Here, we address this critical gap in knowledge and demonstrate that SARS-CoV-2-infected cells escape killing by healthy NK cells in a cell-intrinsic manner, resulting in preferential killing of uninfected bystander cells. The ability of infected cells to evade NK cell recognition requires infection to proceed long enough to allow an infected cell to express SARS-CoV-2 encoded proteins. We demonstrate that this escape mechanism is driven by downregulation of ligands for NKG2D, a critical activating receptor on NK cells. Consistent with our findings, Fielding et al. recently reported that NKG2D-L are downregulated on SARS-CoV-2 infected cells, and find that antibody-mediated killing could overcome this evasion mechanism (Fielding *et al*., 2022). We further demonstrate that this ligand downregulation is driven by the SARS-CoV-2 Nsp1 protein, and show that Nsp1 alone is sufficient to mediate direct NK cell evasion. This has important implications for NK cell-mediated control of SARS-CoV-2, as preferential escape of infected cells with killing of bystander cells could contribute to SARS-CoV-2 pathogenesis.

These results illustrate the importance of examining the temporal dynamics of the NK cell response to SARS-CoV-2-infected cells. Other studies have assessed the ability of NK cells to suppress viral load by co-culturing NK cells with SARS-CoV-2-infected targets immediately after infection; their results suggest that under these conditions, NK cells can at least partially control viral replication (Krämer *et al*., 2021; Witkowski *et al*., 2021; Hammer *et al*., 2022). However, our observations demonstrate that NK cells are no longer able to effectively kill infected cells when added to the culture at later time points following infection, after the expression of viral proteins that suppress the innate immune response. The preferential killing of NKG2D-L-positive bystander cells may have important implications for lung pathology during COVID-19. NKG2D-L can be expressed by most cell types (Lanier, 2015) and are upregulated during viral infections, including HIV (Ward *et al*., 2009) and RSV (Zdrenghea *et al*., 2012), in response to stress (Borchers *et al*., 2006). Therefore, NK cells may actually cause damage to the healthy tissue surrounding infected cells rather than clearing the infection. Our data imply that the timing of NK cell trafficking to the site of infection may be critical in determining whether NK cells are protective or pathogenic in COVID-19, as there is a very narrow window for killing of infected cells before bystander killing could ensue.

Our novel finding that the SARS-CoV-2 protein Nsp1 mediates evasion of NK cell killing has significant implications for both the study of the immune response to coronaviruses and the development of therapeutics for COVID-19. Nsp1 is highly conserved across coronaviruses and is an essential virulence factor; it has been shown to inhibit translation of host antiviral factors across multiple beta-coronaviruses (Kamitani *et al*., 2006; Züst *et al*., 2007; Narayanan *et al*., 2008; Min *et al*., 2020; Schubert *et al*., 2020; Vazquez *et al*., 2021; Yuan *et al*., 2021). One study found that, among nearly 50,000 SARS-CoV-2 sequences analyzed, only 2.4% had any mutations within Nsp1 (Min *et al*., 2020). SARS-CoV-2 Nsp1 also shares 84.4% of its sequence identity with SARS-CoV Nsp1. Moreover, critical motifs within Nsp1 involved in the inhibition of innate immune responses are highly conserved across many beta-coronaviruses(Min *et al*., 2020). On a practical level, the high degree of conservation of Nsp1 and its importance in coronavirus virulence have already made this protein the focus of several therapeutic strategies (Züst *et al*., 2007; Afsar *et al*., 2022; Vora *et al*., 2022). Our work demonstrates that Nsp1 is an even more attractive target than previously thought, as inhibiting the function of this protein has the potential to fully or partially rescue the NK cell response to SARS-CoV-2-infected cells.

Although Nsp1 is a global inhibitor of host translation, our data demonstrate that it has an outsized effect on NKG2D-L and MHC class I surface expression compared to that of other ligands for NK cell receptors. This appears to be due to the varying stabilities of the different ligands on the cell surface, rather than explicit specificity of Nsp1 for NKG2D-L or MHC class I. It has been established that NKG2D-L are rapidly turned over on the cell surface and are quickly lost upon treatment with a protein transport inhibitor such as Brefeldin A (Fernández-Messina, Reyburn and Valés-Gómez, 2016; Toledano *et al*., 2018). MHC class I is similarly transient on the cell surface in the presence of translation inhibition, although its stability varies with haplotype and peptide binding (Yarzabek *et al*., 2018). CD54, which was not affected by Nsp1, is highly stable for at least 48 hours, even after treatment with similar inhibitors (Braun *et al*., 1997). Thus, the differential effects of Nsp1 on various ligands for NK cell receptors are likely explained by the varying kinetics of surface turnover.

One of our findings that has been demonstrated by multiple groups is the downregulation of MHC class I upon SARS-CoV-2 infection. The mechanism of this downregulation remains unclear; while we have demonstrated that Nsp1 is responsible for this phenomenon, ORF3a (Arshad *et al*., 2022), ORF7a (Arshad *et al*., 2022), ORF6 (Yoo *et al*., 2021), and ORF8 (Zhang *et al*., 2021) have also been implicated. According to the well-established “missing self” model of NK cell activation (Ljunggren and Kärre, 1990; Kärre, 2008), the downregulation of self-MHC can induce NK cell activation through subsequent lack of inhibitory signaling through the killer cell immunoglobulin-like receptors (KIRs). Therefore, it might be expected that the downregulation of MHC by SARS-CoV-2 would enhance the ability of NK cells to lyse infected cells–precisely the opposite of what we observed in our study. We hypothesize that this can be explained by 1) the relative magnitudes of MHC class I and NKG2D-L downregulation on infected cells and 2) the accepted dogma in the field that missing self alone is not sufficient to cause robust NK cell activation (Vivier *et al*., 2011; Barrow and Colonna, 2019). As a result, we propose that the loss of NKG2D-L is the dominant factor in the NK cell response (or lack thereof) to SARS-CoV-2.

Our study has several limitations. In order to focus on NK cell responses in the respiratory tract, we used A549-ACE2 cells, which are an immortalized, malignant cell line. This could therefore have enhanced NK cell targeting of bystander cells. Additionally, while we demonstrated that Nsp1 was sufficient to confer NK cell escape, we were unable to test whether the absence of Nsp1 rescues NK cell killing because knockout of Nsp1 is lethal to the virus. We also did not fully evaluate why Nsp1 blocks NKG2D-L more effectively than other proteins, but we hypothesize that these proteins are downregulated first as part of the global translation block because they are turned over on the cell surface more quickly and cannot be replaced. Finally, we did not interrogate the ability of every individual SARS-CoV-2 protein to mediate escape from NK cell killing.

This work has significant implications for the ongoing study of COVID-19. Our results deeply interrogate a major flaw in the ability of the immune system to mount a comprehensive immune response to COVID-19. We demonstrate that the timing of the NK cell response to SARS-CoV-2-infected target cells is critical, with NK cells being able to control viral replication early in infection, but not after expression of viral proteins has begun. This should be further interrogated *in vivo* to explore whether the kinetics of NK cell trafficking during COVID-19 affect disease outcome. Finally, we reveal that SARS-CoV-2 protein Nsp1 is a major factor in mediating evasion of NK cell killing. This finding reinforces the attractiveness of Nsp1 as a therapeutic target.

## Methods

### Cell lines

A549-ACE2s were a gift from Ralf Bartenschlager. VeroE6 cells were obtained from ATCC. Both cell lines were confirmed to be mycoplasma-free. A549-ACE2 cultures were replenished after no more than 25 passages to ensure integrity of ACE2 expression.

### Viral stock generation and titration

icSARS-CoV-2/WA-01-mNeonGreen was a kind gift from Dr. Pei-Yong Shi. Virus was passaged twice in VeroE6 cells and titered by plaque assay on VeroE6 cells using Avicel (FMC Biopolymer) overlay. Passage 3 was used for all experiments. The viral stock was deep-sequenced and aligned to reference genomes in GenBank to confirm sequence.

### Infection with SARS-CoV-2

A549-ACE2 cells were plated in a 12-well plate at a density of 0.1e6 cells/mL the day prior to infection. On the day of infection, A549-ACE2s were washed with PBS (ThermoFisher Scientific, Cat. 10010023), placed in DMEM (Life Technologies, Cat. 11885-092) supplemented with 2% FBS (“D2”) and brought into the BSL3 laboratory. The D2 was removed and virus was added at an appropriate MOI (0.5 unless otherwise noted) in 200 uL D2. Mock-infected cells received 200 uL D2 containing no virus. The infected or mock-infected cells were rocked at 37°C for 1 hour, after which time they were washed with PBS to remove unbound virions. Fresh D2 was then added to the cells and they were replaced into a 37°C incubator for 0-48 hours.

### NK cell isolation and activation

NK cells were isolated from cryopreserved healthy donor PBMC using the Miltenyi MACS Human NK Cell Isolation Kit (Miltenyi, Cat. 130-092-657) according to the manufacturer’s instructions. All donors were collected prior to the advent of the COVID-19 pandemic and were therefore naive to SARS-CoV-2. Following isolation, NK cells were transferred to a round-bottom 96-well plate and resuspended in complete RPMI supplemented with 25 ng/mL rhIL-2 (R&D Systems, Cat. 202-IL-010). NK cells were then placed into a 37°C incubator. After 12-16 hours, the NK cells were washed, counted, and plated in a fresh round-bottom 96-well plate at a concentration of 150,000 cells per well in 100 uL.

### Flow cytometry-based killing and NK cell activation assays

On the day of the experiment, SARS-CoV-2-infected or mock-infected A549-ACE2 cells were trypsinized and counted. The cells were washed and transferred to complete RPMI, then added to the NK cell cultures at an E:T ratio of 5:1 for killing assays and 1:2 for NK cell activation assays. Additional wells containing only target cells or only NK cells were included for control purposes. For activation assays, CD107a-PE, Brefeldin A (eBioscience, Cat. 00-4506-51), and Monensin (eBioscience, Cat. 00-4505-51) were added to all wells at the start of co-culture. Once combined, the NK cells and A549-ACE2s were briefly spun down to bring the cells together and replaced in the incubator for 3 hours (killing assays) or 4 hours (NK activation assays) at 37°C. Following the incubation period, the cells were washed with PBS and stained with the eFluor 780 viability dye. For NK cell activation assays, the cells were stained with a panel of markers against surface markers expressed on NK cells. All assays were then fixed in 4% PFA in PBS (EIS, Cat. 15710) for 30 minutes, transferred to BSL2 facilities, washed, and stored overnight in 1% PFA in PBS at 4°C. The following day, activation assay samples were permeabilized (BD Biosciences, Cat. 340973), stained with a panel of intracellular markers, and collected on a Cytek Aurora for analysis. A table of flow cytometry reagents used can be found in Supplementary Table 1.

### Ligand profiling of SARS-CoV-2-infected and mock-infected cells

On the day of the experiment, SARS-CoV-2-infected or mock-infected A549-ACE2 cells were trypsinized and transferred to a round-bottom 96-well plate. The cells were washed with PBS and stained with the eFluor 780 viability dye. The cells were then stained with a panel of markers against the ligands for 6 different receptors expressed by NK cells (Supp. Table 1). They were then fixed in 4% PFA for 30 minutes, transferred to BSL2 facilities, washed, and stored overnight in 1% PFA in PBS at 4°C. The following day, samples were washed and collected on a Cytek Aurora for analysis.

### RT-qPCR

RNA was extracted cells lysed in DNA/RNA Shield using RNA Clean & Concentrator kits (Zymo Research, Cat. R1018) and excess DNA was removed from the samples using the TURBO DNA-free Kit according to the manufacturer’s instructions (Fisher Scientific, Cat. AM1907). RT-qPCR reactions were prepared using the Invitrogen superscript III Platinum One Step qRT PCR Kit with ROX (Invitrogen, Cat. 11745500) and primer/probe Taqman assays ordered from Thermo Scientific (Supp. Table 2). The QuantStudio 3 Real-Time PCR System was used to quantify transcript levels (Thermo Fisher, Cat. A28567). Three technical replicates of each sample were measured, and all samples were normalized to an endogenous control (*18S*).

### SARS-CoV-2 protein plasmids

Plasmids encoding individual SARS-CoV-2 proteins and GFP were obtained from the Qi lab at Stanford University. Each plasmid included Strep Tag II, allowing for identification of transfected cells that successfully expressed the protein of interest.

### Transient transfection of A549-ACE2s

A549-ACE2s were plated the day before transfection in 24-well plates at a concentration of 75,000 cells per well. Plasmids were transfected into cells with the aid of FugeneHD (Promega, Cat. E2311) using a ratio of 4 uL FugeneHD per 1 ug of plasmid DNA. Four technical replicates of each transfection were performed. The cells were then placed into a 37°C incubator for 48 hours.

### Ligand profiling of transfected cells

48 hours after transfection, transfected A549-ACE2s were harvested and transferred to a 96-well plate for flow cytometry staining. Cells were stained with the eFluor 780 viability dye and a panel of fluorescent antibodies against NKG2D-L, DNAM-1-L, CD54, and MHC class I (Supp. Table 2) before being fixed in 4% PFA for 15 minutes. Fixed cells were permeabilized and stained with a primary antibody against Strep Tag II, washed, and stained with a fluorescent secondary antibody against the primary antibody. Cells were then analyzed on a CyTek Aurora.

### Brefeldin A treatment of A549-ACE2s

A549-ACE2s were cultured in D10 alone or D10 supplemented with either 0.5x or 1x Brefeldin A (eBioscience, Cat. 00-4506-51) for 24 or 48 hours and expression of NK cell receptor ligands was expressed by spectral cytometry.

### ELISA quantification of soluble MICA and soluble ULBP-2

Supernatants from mock or SARS-CoV-2 infected A549-ACE2s were harvested 48 hours post-infection. Triton-X 100 was added to the supernatants to a final concentration of 1% for inactivation of virus and samples were stored at -80°C until use. ELISAs were performed using the Human MICA DuoSet ELISA (R&D Systems, Cat. DY1800) and Human ULBP-2 DuoSet ELISA (R&D Systems, Cat. DY1298) kits according to the manufacturer’s instructions. 1% Triton-X 100 (Sigma-Aldrich, Cat. T9284-100ML) was added to the standards (prior to serial dilution) to account for effects of the inactivation reagent on soluble protein concentration.

### Transfected cell killing assay

IL-2-activated NK cells were co-cultured for 3 hours at an E:T ratio of 5:1 with A549-ACE2s that had been transfected 48 hours earlier with either Nsp1, Nsp14, or GFP. Following the incubation period, the cells were washed with PBS and stained with the eFluor 780 viability dye and an antibody against surface NKG2D-L before being fixed in 4% PFA for 15 minutes. Fixed cells were permeabilized and stained with a primary antibody against Strep Tag II, washed, and stained with a fluorescent secondary antibody against the primary antibody. Samples were then analyzed on a CyTek Aurora.

### Data analysis

Flow cytometry data analysis was performed using FlowJo v10.7.1. Figures were generated in R using the *ggplot2* package. Colors for figures were generated using the *tayloRswift* package. Statistical analyses were performed using the R *ggpubr* package.

## Supporting information

all supplemental materials

## ACKNOWLEDGEMENTS

We are grateful to the donors who provided healthy peripheral blood for these experiments. We thank Dr. Pei-Yong Shi for the kind gift of icSARS-CoV-2/WA-01-mNeonGreen. We thank Aaron J. Wilk for input on manuscript structure. We thank Drs. Rich Stanton and Hugh Reyburn for insightful conversations on the NK cell response to SARS-CoV-2. We thank Dr. Jaishree Garhyan for her assistance in Stanford’s BSL3 facilities. Figure illustrations were created using BioRender.com. Data analysis and visualization was performed using R.

## AUTHOR CONTRIBUTIONS

MJL and CAB conceived the project and designed experiments. MJL, ML, AR, and AB performed experiments. MJL performed statistical analyses under the supervision of SPH and generated figures. AR, LZ, LQ, and SPH provided intellectual input. MJL and CAB wrote the manuscript. All authors reviewed and revised the manuscript.

