## Supplementary material for "SARS-CoV-2 escapes direct NK cell killing through Nsp1-mediated downregulation of ligands for NKG2D": all supplemental materials

### SUPPLEMENTAL DATA

#### Figure S1: NK cells robustly kill uninfected A549-ACE2s in the presence of IL-2 and NK cell subset frequency is not altered upon co-culture

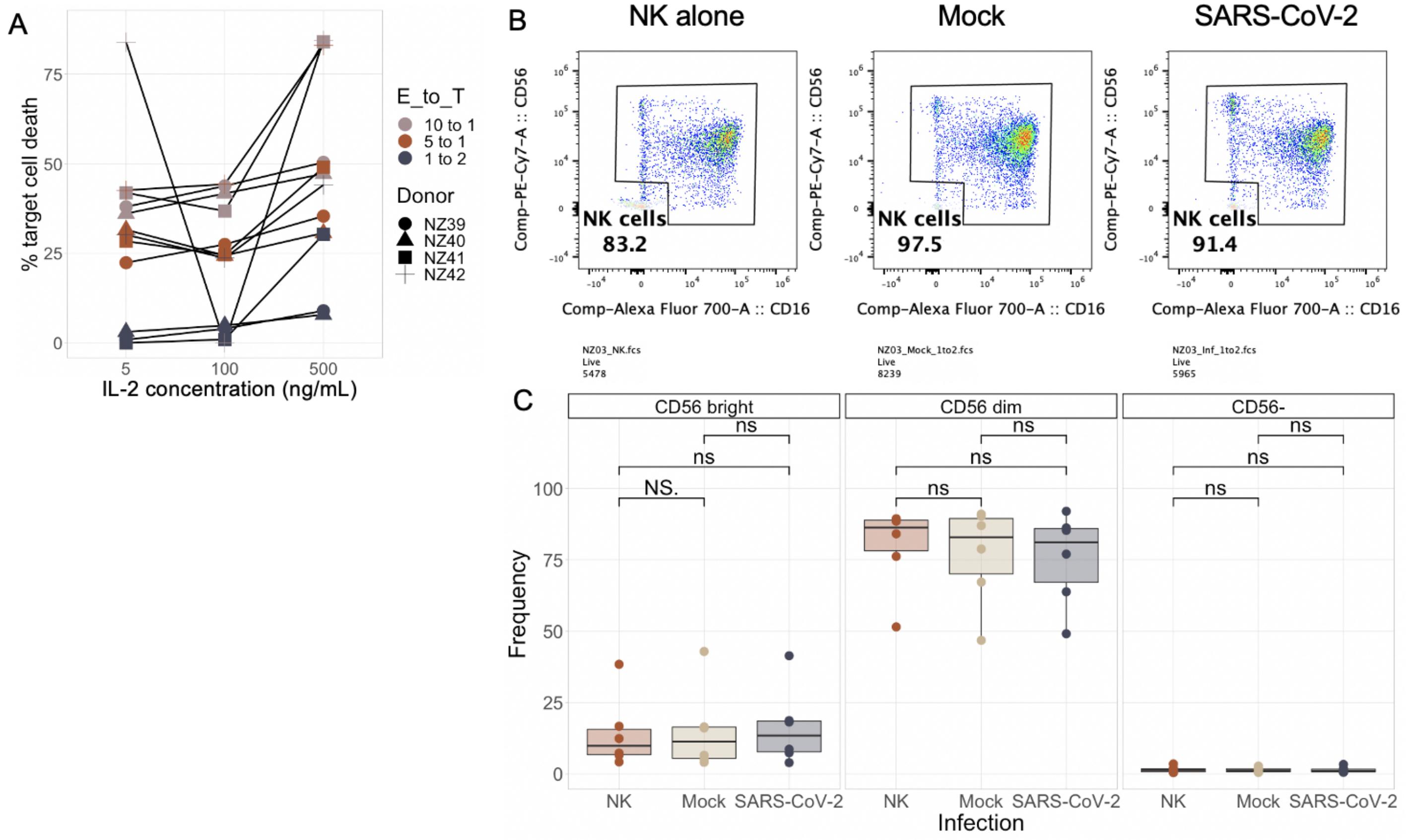


**Figure S1: NK cells robustly kill uninfected A549-ACE2s in the presence of IL-2 and NK cell subset frequency is not altered upon co-culture.** A) Background-subtracted target cell death among A549-ACE2s co-cultured with healthy NK cells for 3 hours. NK cells were preactivated overnight with either 5, 100, or 500 IU/mL rhIL-2 for 12-16 hours. Results are shown at E:T ratios of 1:2, 5:1, and 10:1. B) Representative flow plots showing CD56 and CD16 expression in NK cells alone or co-cultured with uninfected or SARS-CoV-2-infected A549-ACE2s. C) Boxplots showing the frequencies of $CD56^{bright}$, $CD56^{dim}$, and $CD56^{-}$ NK cells as a percentage of all NK cells across infection conditions.

### Figure S2: Expression of NK receptor ligands in SARS-CoV-2 infection

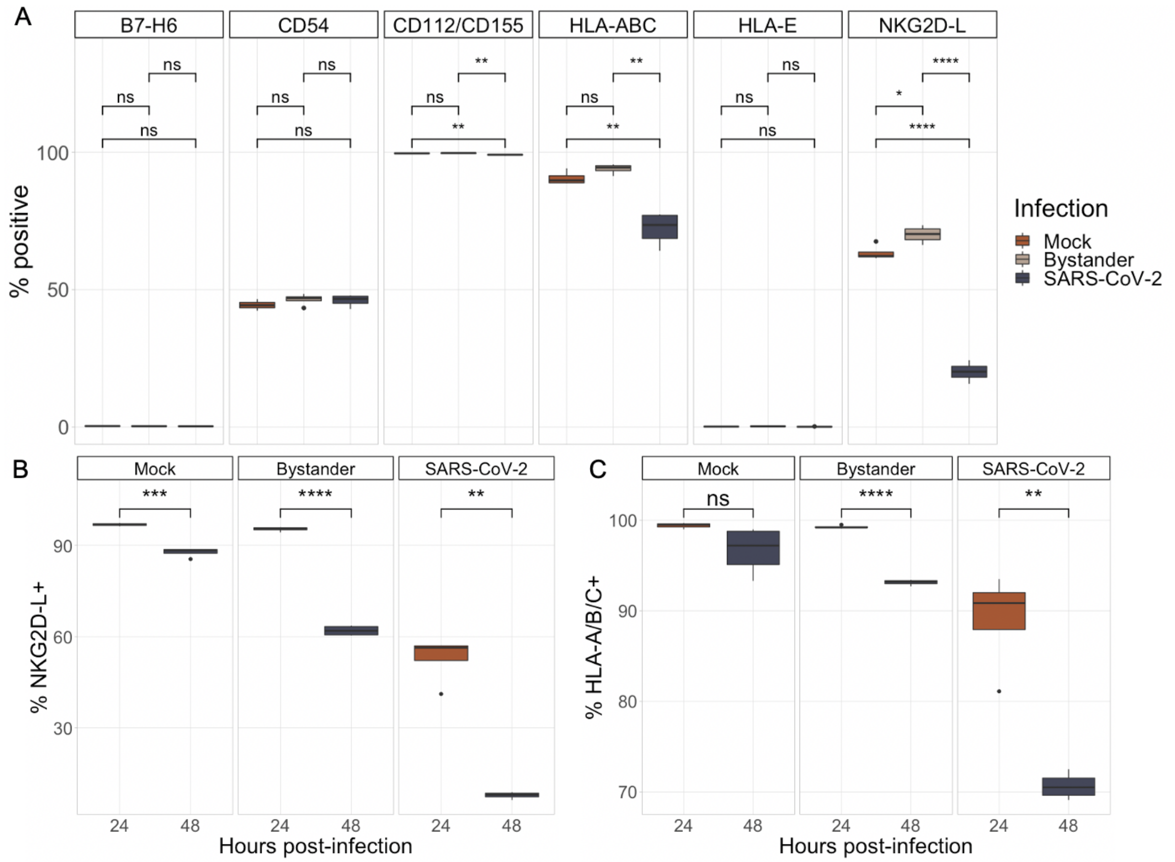


**Figure S2: Expression of NK receptor ligands in SARS-CoV-2 infection.** A) Boxplots showing the percentage of A549-ACE2s expressing the ligands for various NK cell receptors across infection conditions at 48 hours post-infection. Results are shown from an experiment in which cells were infected at an MOI of <u>0.05</u>. B) Percentage of A549-ACE2s expressing NKG2D-L at 24 vs 48 hours post-infection across infection conditions. C) Percentage of A549-ACE2s expressing HLA-A/B/C at 24 vs 48 hours across infection conditions. All boxplots represent n=4 technical replicates.

**Figure S3: Validating identification of transfected cells**

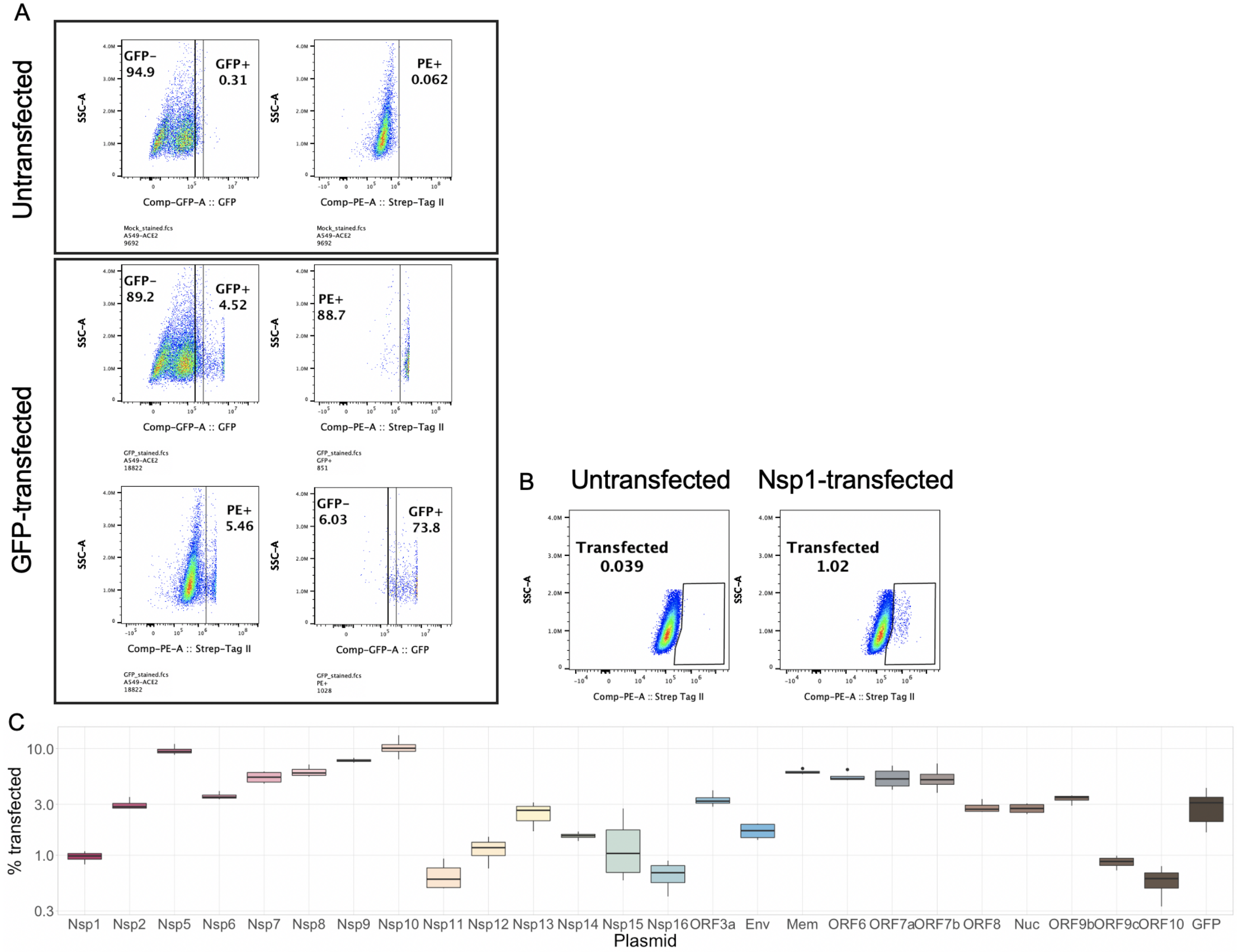


**Figure S3: Validating identification of transfected cells.** A) Representative flow plots showing expression of GFP and Strep Tag II in mock-transfected versus GFP-transfected A549-ACE2s to illustrate overlap between GFP expression and Strep Tag II expression. B) Representative flow plots of Strep Tag II expression in mock-transfected vs Nsp1-transfected A549-ACE2s. C) Boxplots showing the percentage of cells positive for Strep Tag II by flow cytometry following transfection with plasmids encoding various proteins. All boxplots represent n=4 technical replicates for all plasmids.

### Figure S4: Frequency of CD54- and DNAM-1-L-expressing cells after transfection of SARS-CoV-2 proteins

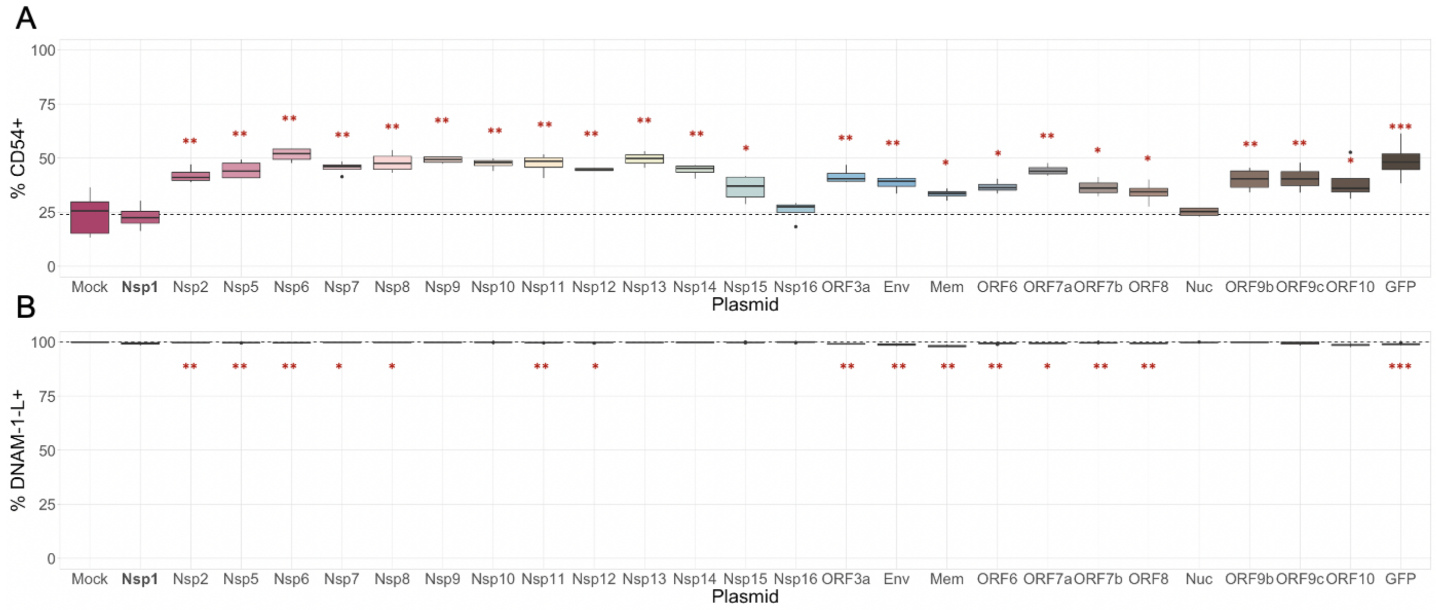


**Figure S4: Frequency of CD54- and DNAM-1-L-expressing cells after transfection of SARS-CoV-2 proteins.** A-B) Percentage of A549-ACE2s expressing CD54 (A) or CD112/CD155 (B) by flow cytometry following transfection with plasmids encoding various proteins. Dashed lines represent the mean of mock-transfected samples expressing the ligand of interest. All boxplots represent n=4 technical replicates for all plasmids and n=8 for all "mock" samples.

### Figure S5: ELISA data for detection of soluble MICA (sMICA) and soluble ULBP-2 (sULBP2)

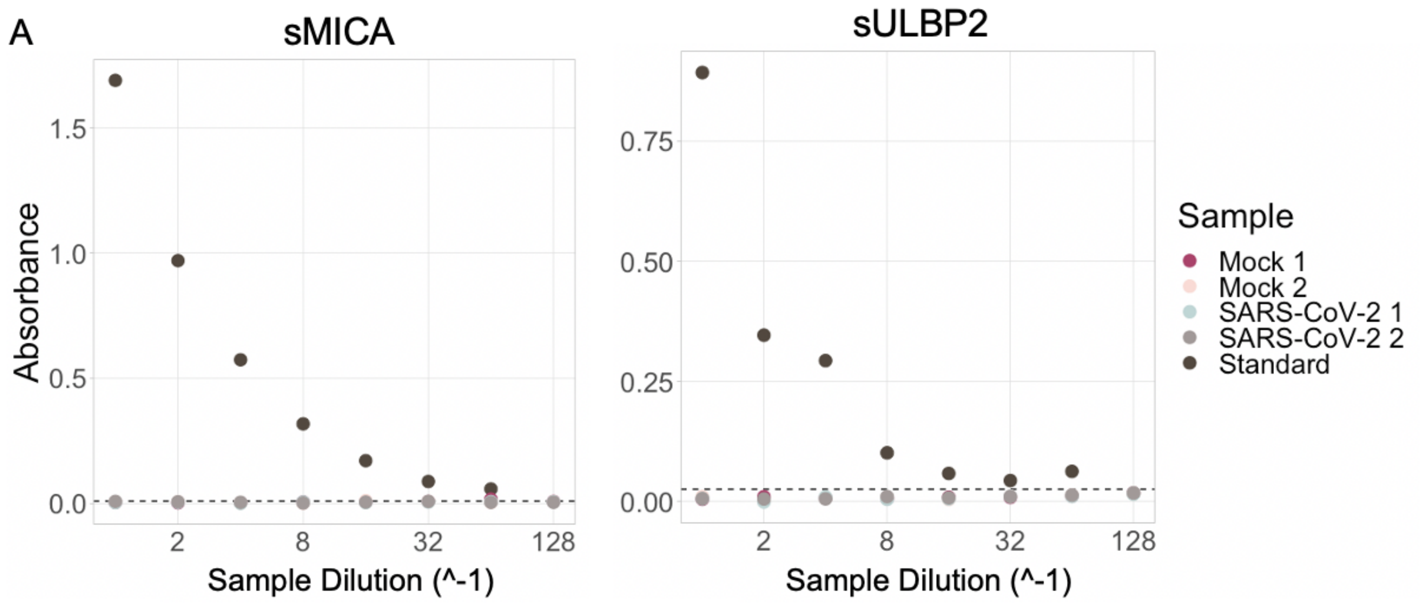


**Figure S5: ELISA data for detection of soluble MICA (sMICA) and soluble ULBP-2 (sULBP2).** A) Absorbance values of the supernatants from mock or SARS-CoV-2-infected cultures at varying dilutions as measured by plate-based ELISAs for soluble MICA (sMICA) and soluble ULBP2 (sUPBP2). Absorbance values were calculated by subtracting absorbance readings taken at 560 nm from those taken at 450 in accordance with the manufacturer's instructions. Dashed lines indicate limits of detection.

### Figure S6: Effects of Brefeldin-A treatment on A549-ACE2 NK receptor ligand expression

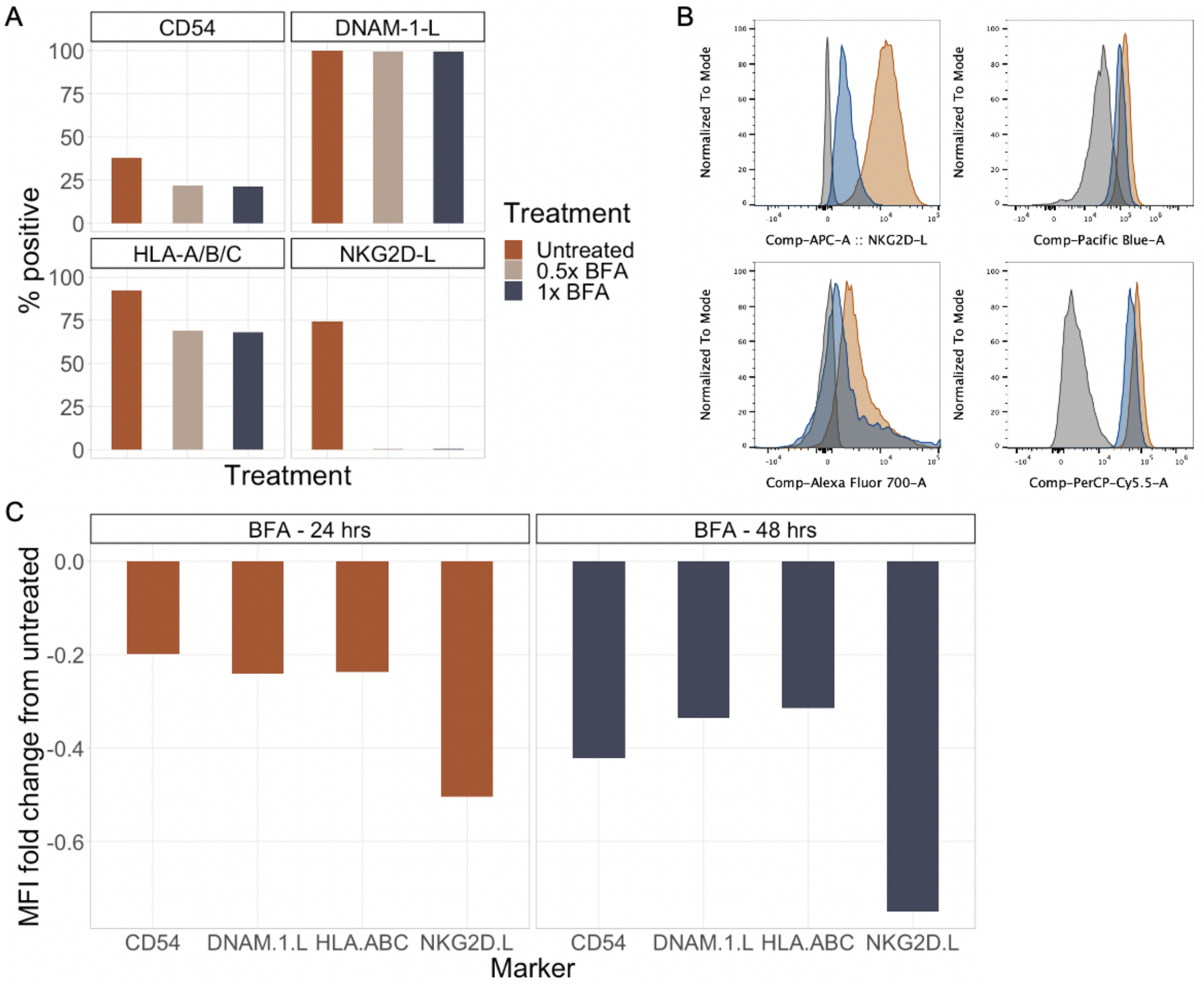


**Figure S6: Effects of Brefeldin-A treatment on A549-ACE2 NK receptor ligand expression.** A549-ACE2s were cultured for 48 hours with Brefeldin A (1x or 0.5x) or media alone. A) Percentage of A549-ACE2s expressing CD54, DNAM-1-L (CD112/CD155), HLA-A/B/C, or NKG2D-L after 48 hours under each treatment condition. Bars represent the mean of n=2 technical replicates. B) Representative histograms showing expression of these markers in BFA (1x)-treated or untreated A549-ACE2 after 48 hours. C) Bar plots showing the fold changes in MFI of each ligand on A549-ACE2s treated with 1x BFA compared to untreated cells after 24 (left) or 48 (right) hours. Bars represent the mean of n=2 technical replicates.

### Figure S7: Identification of transfected cells in transfected cell killing assays

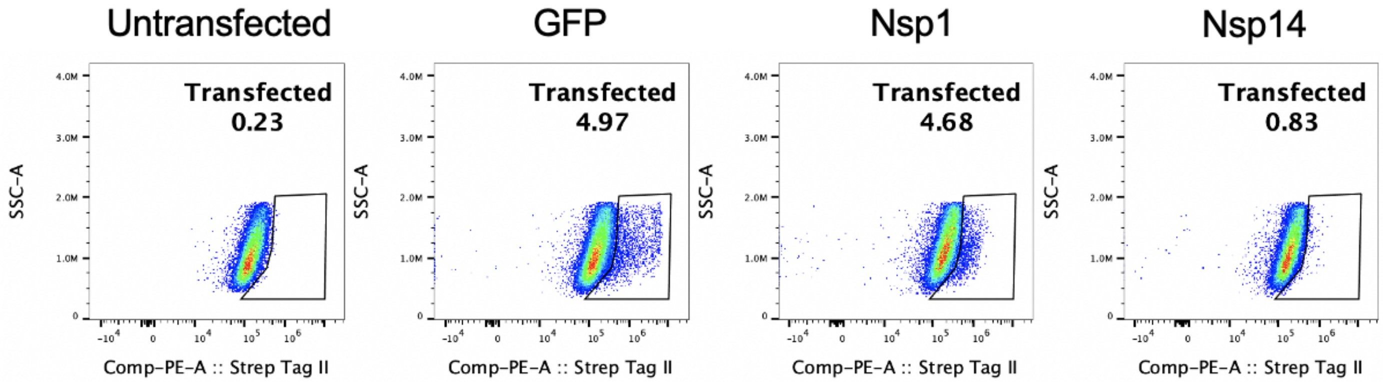


**Figure S7: Identification of transfected cells in transfected cell killing assays.** Representative flow plots showing the percentage of transfected A549-ACE2s upon transfection with various plasmids.

**Supplementary Table 1: Flow cytometry reagents**

| Antigen | Fluor | Clone | Manufacturer |
|---|---|---|---|
| CD56 | PE-Cy7 | HCD56 | Biolegend |
| CD16 | Alexa Fluor 700 | 3G8 | Biolegend |
| DNAM-1 | APC | 11A8 | Biolegend |
| NKG2D | BV785 | 1D11 | Biolegend |
| CD38 | BV605 | HIT2 | Biolegend |
| CD69 | PE-Dazzle594 | FN50 | Biolegend |
| CD107a | PE | H4A3 | Biolegend |
| HLA-E | PE-Cy7 | 3D12 | Biolegend |
| B7-H6 | PE | 875001 | R&D Systems |
| CD112 | PerCP-Cy5.5 | TX31 | Biolegend |
| CD155 | PerCP-Cy5.5 | SKII.4 | Biolegend |
| MICA | APC | 159227 | R&D Systems |
| MICB | APC | 236511 | R&D Systems |
| ULBP-1 | APC | 170818 | R&D Systems |
| ULBP-2,5,6 | APC | 165903 | R&D Systems |
| CD54 | Alexa Fluor 700 | HA58 | Biolegend |
| HLA-A/B/C | Pacific Blue | W6/32 | Biolegend |
| Live/Dead | eFluor 780 | N/A | ThermoFisher |
| Strep Tag II | N/A | ab76949 | Abcam |
| Anti-rabbit IgG | PE | Poly 4064 | Biolegend |

**Table S1: Flow cytometry reagents.** Relevant information regarding all antibodies and dyes used for flow cytometry experiments described in this manuscript.

**Supplementary Table 2: RT-qPCR reagents**

| Gene | Assay ID / Cat # | Manufacturer |
|---|---|---|
| *MICA* | Hs07292198_gH | ThermoFisher Scientific |
| *MICB* | Hs00792952_m1 | ThermoFisher Scientific |
| *ULBP1* | Hs0036941_m1 | ThermoFisher Scientific |
| *ULBP2* | Hs01127964_m1 | ThermoFisher Scientific |
| *SARS-CoV-2 N* fwd primer | nCoV-N1-F-100 | Biosearch Technologies |
| *SARS-CoV-2 N* rev primer | nCoV-N1-R-100 | Biosearch Technologies |
| *SARS-CoV-2 N* probe | nCoV-N1-P-25 | Biosearch Technologies |
| *18S* | 4352930E | ThermoFisher Scientific |

**Table S2: RT-qPCR reagents.** Relevant information regarding all primers and probes used for RT-qPCR experiments described in this manuscript.
